# Successful reproduction of *Trachemys scripta* in the Altrhein of Kehl (Germany) and simultaneous increase of estimated population size

**DOI:** 10.1101/2020.08.06.240788

**Authors:** Carsten Schradin

## Abstract

The European Union categorised pond sliders (*Trachemys scripta*) as an invasive species for which all member countries have to develop an action plan. While *T. scripta* has been released illegally in thousands of freshwaters in Europe, monitoring programs in Germany are missing. Based on short term surveys, anecdotal data, and assumptions, the action plan in Germany proposes that *T. scripta* does not represent a problem as the cold weather causes high mortality and does not allow for successful reproduction. Here I present data of five years of monitoring at the Altrhein of Kehl, at the border between Germany and France showing: 1. Apart from *T. scripta*, five additional species have been released (*Pseudemys nelson, Pseudemys concinna, Graptemys pseudogeographica, Mauremys spec*., *Chrysemys picta*.). 2. Population size seemed to increase over the years, with 116 individuals being counted in 2020 compared to 33 individuals in 2016. 3. Successful reproduction of. *T. scripta* was indicated by the observation of very small individuals and proved by the collection of hatchlings in two years. While the data presented here would benefit from more years of more intensive monitoring, they are sufficient to draw the conclusion that the German action plan is insufficient because individuals survive for long periods and do reproduce successfully. Similar results have been reported from other central European countries. Thus, *T. scripta* might become invasive not only in south but also in central Europe. In sum, national action plans of how to deal with the invasive *T. scripta* should to be revised in Germany and many other European countries.

## INTRODUCTION

Invasive species are threatening native biodiversity worldwide. Neobiota are alien species that establish themselves outside their natural distribution range due to human action, i.e. they reproduce successfully in the areas in which they have been released and they maintain their population (Geiger & Waitzmann 1996; Wilson et al. 2009). They are regarded as being invasive if they further spread their distribution into new areas and at the same time cause harm on the native fauna and / or flora by predation and competition (Lowe et al. 2000; Wilson et al. 2009).

From the worlds’s 100 worst invasive alien species, only two species are reptiles, the brown tree snake (*Boiga irregularis*) from Australia and the North American red-eared slider (*Trachemys scripta*) (Lowe et al. 2000). Compared to invertebrates (26 of the 100 worst invasive species) and small mammals (10 of 100 the worst invasive species), reptiles grow, mature, and reproduce slowly, which can be one reason why so few reptiles are invasive. However, a slow life history also makes it more difficult to monitor and predict the long-term consequences of an introduced alien species, i.e. whether it will establish itself and become invasive or not. From an ecological point of view it is not important whether an invasion takes a few years or 100 years, as the long-term consequences on the native flora and fauna would still be deleterious. The potentially invasive *T. scripta* has been released worldwide, including central and south America (Böhm 2013), Asia (Mo 2019), and Europe (Standfuss et al. 2016). While the slow life history of the species would allow sufficient time for authorities to react in time, unfortunately the opposite occurs. Small neobiota populations are often ignored by the authorities, and long-term monitoring programs are only established many years after the population has grown to a size that makes population control difficult (Sancho & Lacomba 2016). While *T. scripta* has become invasive in areas with tropical, sub-tropical, and Mediterranean climate (Lowe *et al*. 2000; Foglini & Salvi 2017), the consequences in areas with temperate climate are so far not clear (Cadi et al. 2004; Prevot et al. 2007; Kopecký, Kalous & Patoka 2013; Standfuss et al. 2016).

In 2014, the European Union published Regulation (EU) No 1143/2014, declaring an action plan to prohibit the import, breeding and releasing of invasive species (European_Parliament & Council_of_the_European_Union 2014), including *T. scripta* (European_Commission 2016). A regulation is a legal act of the European Union that becomes immediately enforceable as law in all member states. Thus, this regulation forces all member states to take actions to avoid the spread of the declared invasive species. As the environmental conditions such as flora, fauna, and climate differ between members states, the individual states are allowed to respond differently. In Germany, the proposed actions against *T. scripta* are to increase public awareness, allowing non-commercial transfer of individuals that are already kept in captivity, while the removal of individuals from wild habitats is regarded to be very costly and ineffective and thus in most circumstances not feasible (StA_"Arten-_und_Biotopschutz” 2018a). Instead, in Germany it is assumed that due to the cold weather mortality of released *T. scripta* is high (Geiger & Waitzmann 1996) and that successful reproduction only occurs sporadically (Pieh & Laufer 2006) such that populations are not stable (StA_"Arten-_und_Biotopschutz” 2018a).

The aim of this study was to monitor a known population of *T. scripta* at the Altrhein of Kehl in Germany (Pieh & Laufer 2006; Laufer 2007) to determine 1. if there is evidence of population decrease or increase, 2. whether there is evidence of successful reproduction of this population, and 3. whether there is evidence that additional individuals are released into the wild. As such, the overall aim was to establish one of the first long-term monitoring programs in Germany to obtain information on the population dynamics of the alien and potentially invasive species *T. scripta*.

## MATERIALS AND METHODS

### Study area and study period

The study was conducted in the city of Kehl in Baden-Württemberg, Germany. Kehl is at the Upper Rhine Valley close to the French city of Strasbourg. Data were collected at the Altrhein of Kehl, which was a part of the river Rhein more than 100 years ago. It now represents a pond, 690m long and between 25 and 80m wide. The climate in Kehl is temperate, but the Upper Rhine Valley is one of the warmest areas of Germany. During the study period, the minimum temperature in winter was around -12°C (2016), and daily maximum temperatures in summer were around 40°C in 2019. Periods of permanent frost in winter were relatively short with 1-2 weeks in the years 2016-2019, and no such period in 2020.

The study was conducted from May to July in the years 2016 to 2019 as part of a course I gave at the Hector Akademie Kehl. In this course, I taught highly gifted school children 8-9 years old about ecology and nature conservation. Due to the Corona crisis, this course was cancelled in 2020. In this year, I used the opportunity to collect data on two sunny days in March (at this stage it was unsure whether later data collection would be possible) as well as during 6 afternoons in May and June. There was no indication that the March data differed from the later data (highest number of individuals and species was recorded in May and June, the normal period of monitoring), and excluding these data would not have changed the results.

### Monitoring

In the years 2016-2018, the population of exotic pond turtles was monitored during six afternoons, in 2019 during seven, and in 2020 during eight. During the monitoring, I walked along the East coast of the Altrhein and noted any pond turtle observed. On five locations that were previously determined to have a high abundance of pond turtles, I stopped and watched the area carefully with binoculars, but also any pond turtle observed between these spots was recorded. For every individual, the size was estimated with the help of a template. I used the following categories for estimating the diameter of the carapax: 5cm, 10cm, 20cm or 30cm.

In 2016, only the number and size of observed pond turtles was noted. In this year it became evident that not – as originally expected - only individuals of the species *Trachemys scripta* were present, but also several other species. In the following years, I took photographs of each of the individual I observed. The distance to the observed pond turtles were up to 30m. From 2017 to 2019 photographs were taken with a Panasonic HC-V380EG-K camcorder with 50x optical zoom, and in 2020 with a Nikon Coolpix P1000 Digital camera with 125x optical zoom. Using characteristics of the carapax and especially the head, the genus, species and if possible sub-species of the photographed individuals was determined (Pellerin 2014). If the individual was too far away, in a bad angle such that I could not photograph the head, or partly hidden, the genus could not be identified and the individual was scored as “unknown genus”.

### Measurement of hatchlings

Each year as part of a scientific outreach, I held an information afternoon with children between the ages 5-6 years old, and their teachers from a Kindergarten next to the Altrhein. The teachers informed me about previous encounters with pond turtles at the kindergarten. They contacted me repeatedly in 2019 and 2020, when hatchlings of *T. scripta* were found at the kindergarten, which I could measure and then send to an animal shelter in Munich (Germany) that specialises in reptiles (https://www.reptilienauffangstation.de/). I was also informed about sightings of large females that were believed to have left the Altrhein to lay eggs.

### Data analysis

Statistical tests were run with GraphPad Instat 3.05. The minimum number alive was determined as the largest number of pond turtle observed during any afternoon per year. To determine the minimum number of individuals per genus / species, the same approach was used, but here data from different taxa could have been from different days (for example minimum number alive for *Trachemys scripta* might have been from another day than from *Pseudemys concinna*). The same procedure was used for individuals of different estimated carapax sizes, causing that the total number of individuals for all size categories is somewhat higher than the minimum number alive

## RESULTS

### Monitoring Altrhein

The annual minimum number alive increased continuously from 33 individuals in 2016 to 166 individuals in 2020 (Fig. 1), which resulted in a significant correlation between year and minimum number alive (Spearman rank correlation, r_s_=1.00, N=5, p=0.02). This was in contrast to the previously published expectation (Pieh & Laufer 2006) that the population size should decline from year to year (Fisher’s exact test, p=0.01).

**Figure 1.**
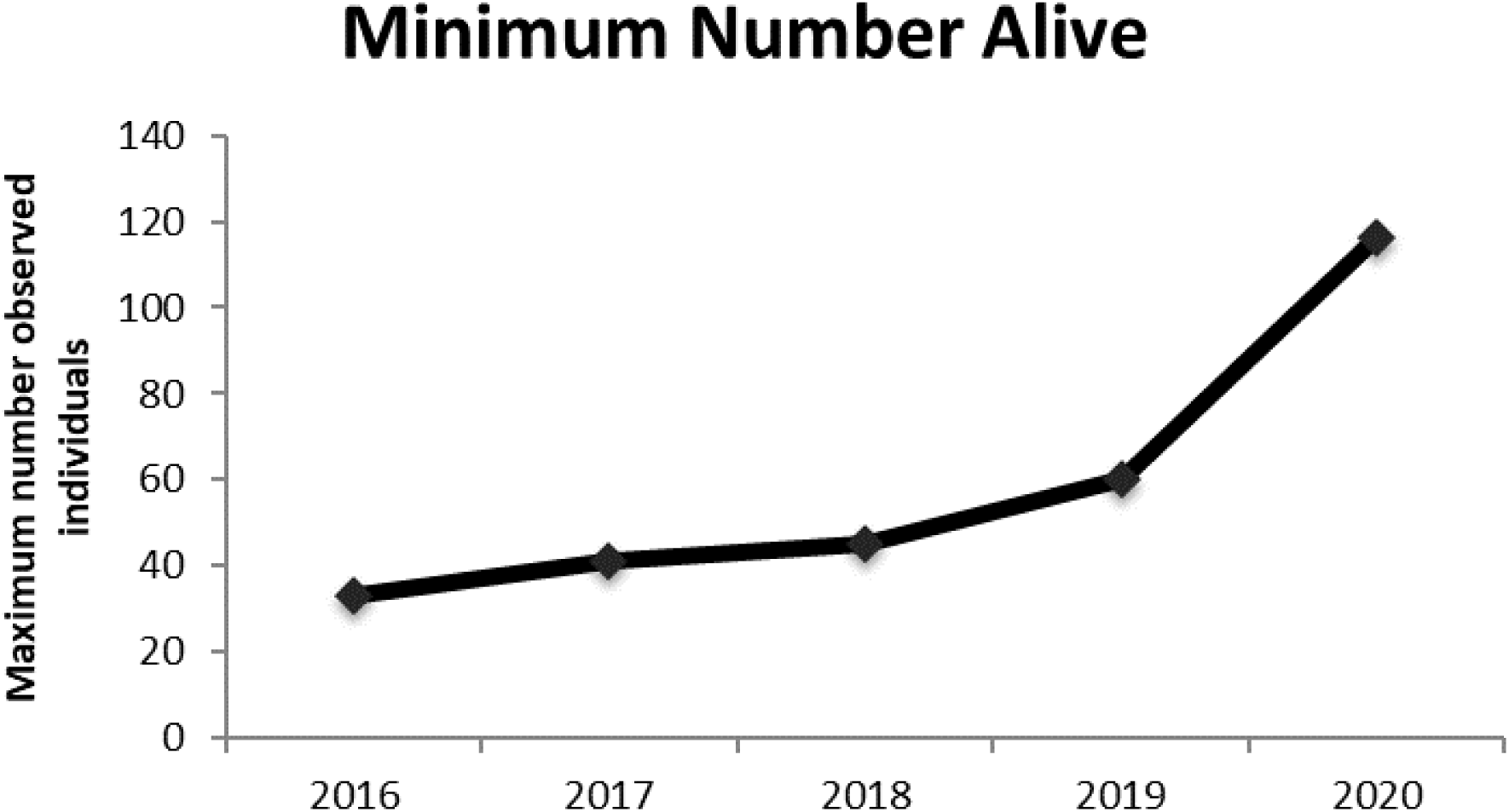
Minimum number alive per year, i.e. the maximum number of individuals observed during one afternoon.

In the four years 2017 to 2020, increasing numbers of very small pond turtles with a carapax of not more than 5cm were observed Spearman rank correlation, r_s_=0.90, N=5, p=0.08, Fig. 2). All photographed pond turtles in this category were of the species *Trachemys scripta*. Additionally, the number of large pond turtles with a carapax of more than 20cm increased over the years (Spearman rank correlation, r_s_=0.90, N=5, p=0.08, Fig. 2). All pond turtles categoriesed as having a carapax of 30cm belonged to one of the three following species: *Trachemys scripta, Pseudemys concinna, Pseudemys nelsoni*.

**Figure 2.**
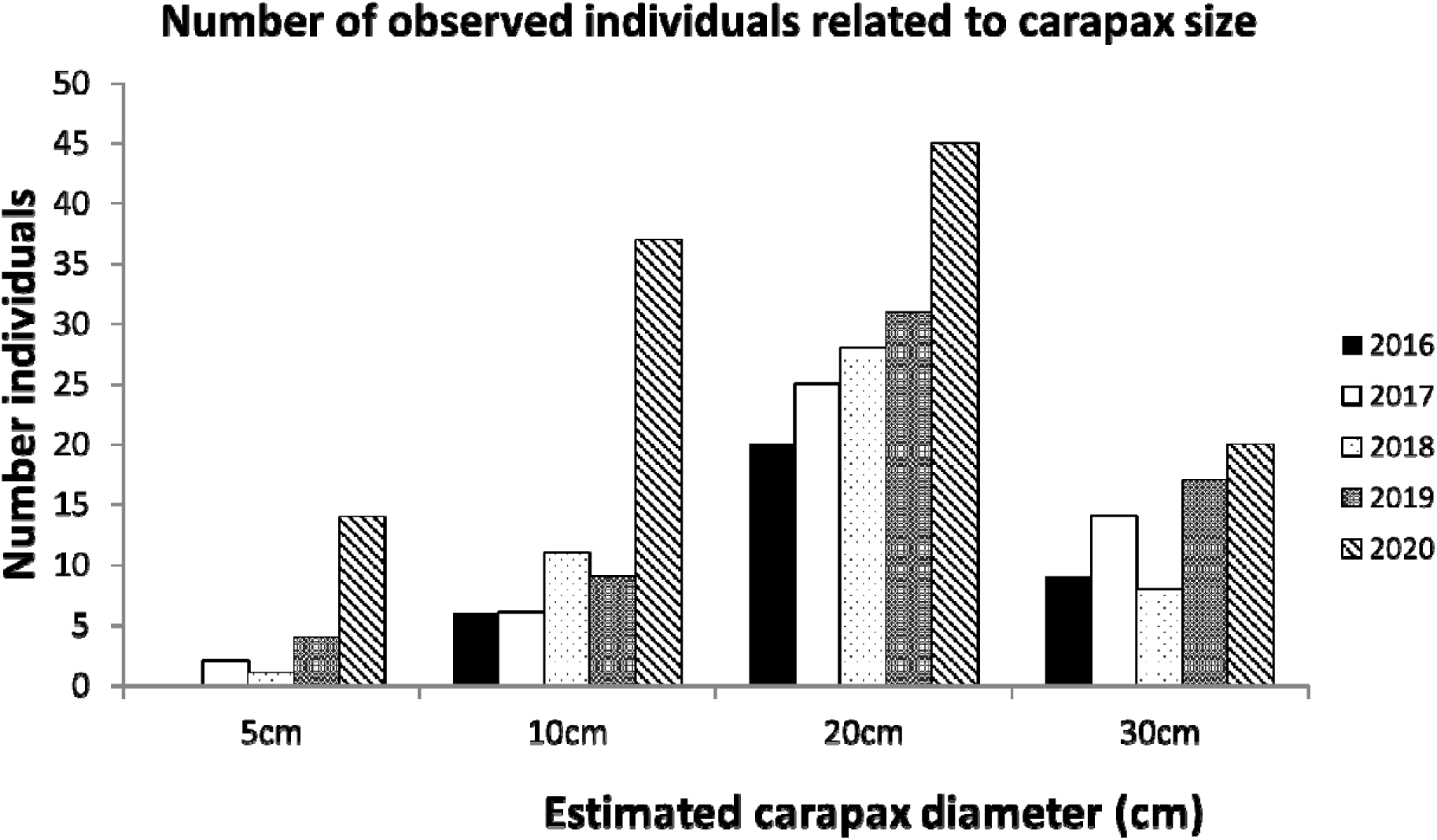
Maximum number of individuals observed during one year related to estimated carapax size. Note that the total number per year is somewhat higher than the minimum number alive, as the numbers reported here are from different days combined.

Altogether 6 different species were observed, with 3 subspecies of *Trachemys scripta* (Table 1). Over the years the number of observed individuals of *T. scripta* seemed to increase (Fig. 3). The two species of *Pseudemys* and *Graptemys pseudogeographica* occurred in all years. Single individuals of *Chrysemys picta* and *Mauremys spec*. (probably *M. sinensis*) were observed in all years except 2018 (Fig. 3).

**Table 1.** Number of individuals observed per taxa per year. Cloudy weather in 2018 made it difficult to determine the correct subspecies / species of both *Trachemys* and *Pseudemys*. The numbers reported here are from different days combined. Individuals for which the taxa could not be determined are excluded, such that the reported numbers here are smaller than the minimum number alive.

| Genus, species, or sub-species | Minimum number 2017 | Minimum number 2018 | Minimum number 2019 | Minimum number 2020 |
| --- | --- | --- | --- | --- |
| *Trachemys scripta* undetermined sub-species | 1 | 7 | 5 | 14 |
| *Trachemys s. elegans* | 8 | 5 | 11 | 24 |
| *Trachemys s. scripta* | 4 | 1 | 6 | 4 |
| *Trachemys s. troosti* | 1 | 0 | 5 | 1 |
| *Pseudemys spec.* | 3 | 12 | 2 | 1 |
| *Pseudemys nelsoni* | 3 | 1 | 1 | 7 |
| *Pseudemys concinna* | 10 | 3 | 7 | 26 |
| *Graptemys pseudempgographica* | 6 | 3 | 4 | 7 |
| *Mauremys spec.* | 1 | 0 | 1 | 2 |
| *Chrysemys picta* | 1 | 0 | 1 | 1 |
| **TOTAL** | **38** | **32** | **43** | **87** |

**Figure 3.**
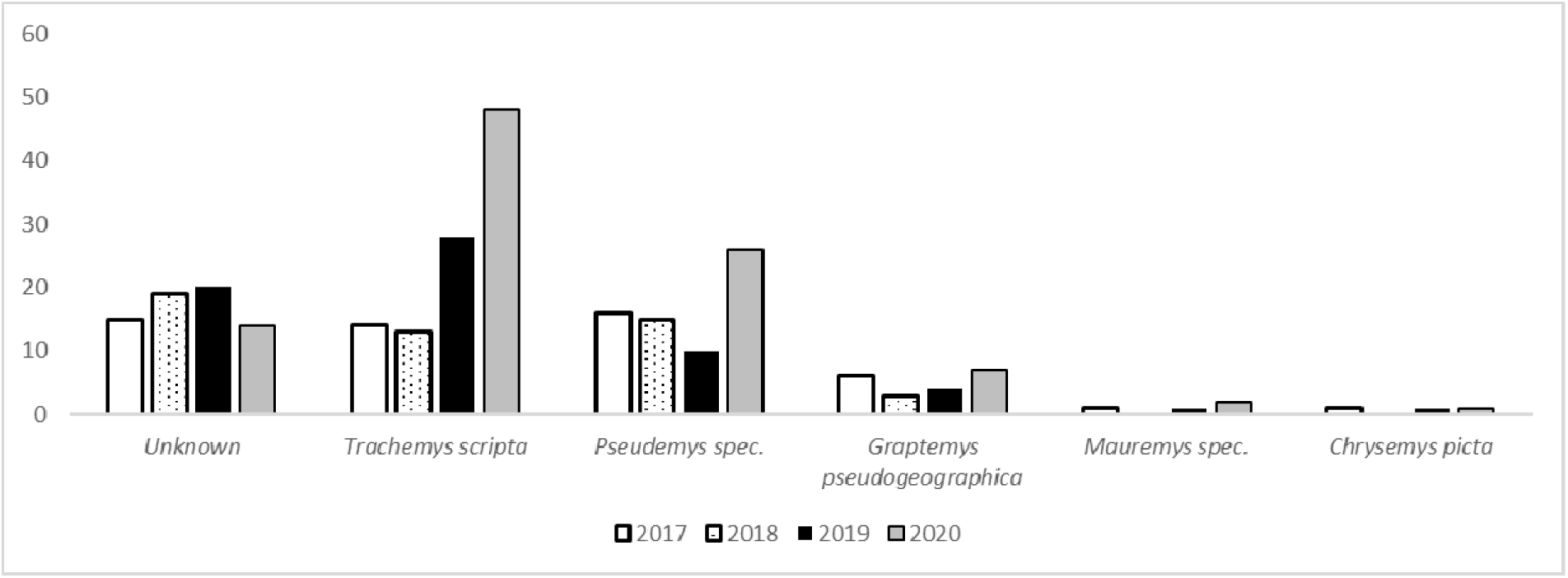
Number of individuals observed per taxa per year. The numbers reported here are from different days combined, such that the reported numbers here can be higher than the minimum number alive.

### Breeding of *Trachemys scripta*

Altogether, 11 very small *T. scripta* were found on land, walking from a park towards the Altrhein: eight in the period 12^th^ to 22^nd^ May 2019, one the 15^th^ of July 2019, and three between the 14^th^ May and 30^th^ June 2020. Two of these were found dead on the bicycle path between the kindergarten and the Altrhein, one died in the aqua-terrarium of the Kindergarten. These dead individuals were disposed of by the kindergarten. The remaining nine individuals were kept and I was able to measure them, confirming that these were all hatchlings of *T. scripta* (Fig. 4, Table 2).

**Table 2.** Measurements of nine young *Trachemys scripta* found at the Altrhein of Kehl, six found in 2019, three in 2020. Measurements in mm if not stated otherwise.

| Trait (mm or g) | Mean | Standard deviation | Maximum | Minimum |
| --- | --- | --- | --- | --- |
| Carapax length | 30.6 | 1.3 | 32.0 | 28.0 |
| Carapax width | 30.4 | 1.3 | 32.0 | 29.0 |
| Carapax height | 15.8 | 1.5 | 19.0 | 13.5 |
| Plastron length | 28.7 | 2.0 | 31.0 | 26.0 |
| Plastron width | 22.3 | 2.1 | 27.0 | 19.5 |
| Body mass (g) | 6.5 | 0.7 | 7.8 | 5.3 |

**Figure 4.**
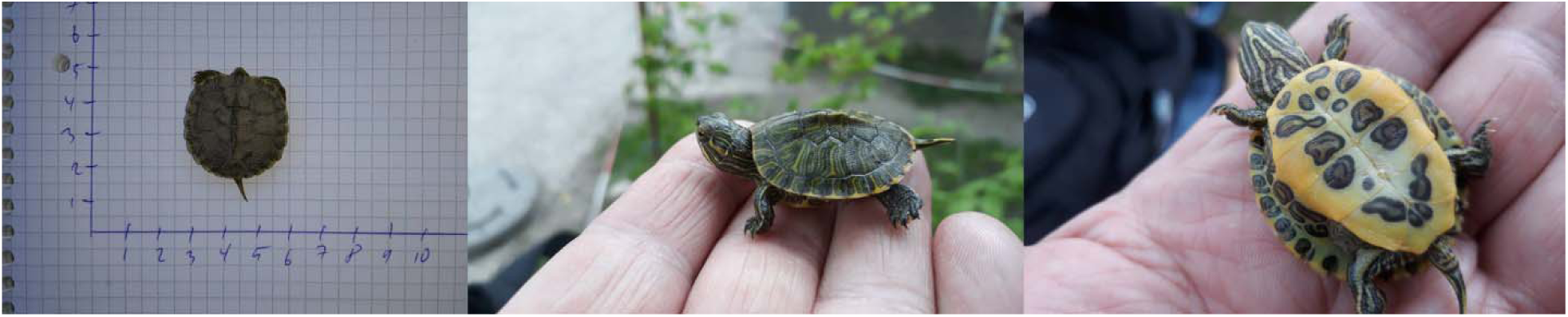
Photos of three different *T. scripta elegans* hatchlings found at the Altrhein of Kehl in May 2019.

The 6^th^ June 2019 a female *T. scripta* was caught at the kindergarten, with a body mass of 3064g and a carapax length of 29cm. A female *T. scripta elegans* was seen laying eggs in the park behind the Kindergarten the 15. June 2020. I could dig out the clutch the 16^th^ of June, which was 15cm below ground. There were a total of 24 eggs with a length of 32.8 ± 1.2mm and a width of 23.9 ± 0.7mm. The total clutch weight was 218g which represented 9.1g per egg.

The 30^th^ of June 2020 a female *Pseudemys concinna* was caught at the kindergarten (and sent by me to Munich the 2^nd^ of July) which weighted 2604g (after it had laid eggs) and had a carapax length of 30cm. In captivity it laid 14 eggs with a length of 36.2 ± 0.5mm and a width of 24.1 ± 1.1mm, and a clutch mass of 166g (11.9g per egg).

## DISCUSSION

In the current study I could show that the population of exotic pond turtles in the Altrhein of Kehl consists of six species, with the invasive species *T. scripta* being the most abundant. In contrast to previous predictions (Pieh & Laufer 2006), there was no indication that the population size declined over the study years. On the contrary, population size seemed to steadily increase, maybe due to the continued release of individuals, but also due to successful reproduction. My results indicate that the assumption is wrong that in Germany released exotic pond turtles will disappear by themselves due to the cold climate.

I expected to observe mainly individuals from the three different subspecies of *T. scripta* in the Altrhein of Kehl and, based on previous reports, maybe some individuals of *Graptemys pseudogeographica* (Pieh & Laufer 2006; Laufer 2007). I was therefore surprised to find a high exotic biodiversity with 4 regular species (*T. scripta, Pseudemys nelsoni, P. concinna*, and *G. pseudogeographica*) and two species (*Mauremys sinensis* and *Chrysemys picta*) with only one individual, but observed in different years (Table 1). This high diversity is most likely a consequence of the trade ban on *T. scripta*, leading to the selling of other species (Maceda-Veiga et al. 2019), which in turn are then released into the wild. Overall, the population size of all pond turtles combined continuously increased over the years (Fig. 1) which was due to an increased number of both small and large individuals (Fig. 2), especially of *T. scripta* (Fig. 3).

I previously pointed out the importance of long-term studies on individually marked individuals (Schradin & Hayes 2017). In contrast, the monitoring methods used in the current study were of low reliability, as no individuals were marked for individual identification, and the study period was relatively short. Nevertheless, the data give no indication that the population declined due to sub-optimal climate as predicted for Germany in general (Geiger & Waitzmann 1996; StA_"Arten-_und_Biotopschutz” 2018a) and for this specific population in particular (Pieh & Laufer 2006). Thus, even though data from more years and from marked individuals would be desirable, this would not affect the current conclusions that (1) there is no indication for a population decline and that (2) *T. scripta* reproduces successfully in Germany. As these conclusions are important for management decisions, it is important not to delay publication of these data.

I observed that the population size increased over the five study years. This could be partly due to the release of more individuals into the wild, based on two points of evidence: Firstly, a high proportion of the observed individuals were from another genus than *Trachemys*, indicating that species that are still available in pet shops are regularly released into the wild. Secondly, a constant increase of the size category of carapax diameter 20cm (Fig. 2), indicates that old individuals have been released, that might have been bought long time ago, and now became too big and as such inconvenient for the pet owners.

In southern France, clutch size varied between 4 and 11 eggs (Cadi et al. 2004), while here I found one clutch consisting of 24 eggs, and observed one very large female of 3kgs that probably laid a clutch. That females lay eggs is not surprising as even without fertilisation females produce eggs. By itself, this does not give evidence for successful reproduction as its neither known whether eggs were fertilised nor whether under the given climatic conditions eggs would hatch.

Very small *T. scripta* individuals have been observed in four out of five study years, making it likely that reproduction occurred repeatedly. The catching of hatchlings in both 2019 and 2020 proved successful reproduction. This is the second case of proven reproduction in Kehl with the first case having been reported for 2004 (Pieh & Laufer 2006). The lacking report of reproduction in Kehl for the years 2005 to 2016 can be explained by a lack of monitoring during these years. The seven hatchlings found in Kehl within 10 days in May plus the single hatchling found in July indicates that these might be from two or three different clutches. The hatchlings probably originate from clutches laid in summer 2018, and might have overwintered in the nest burrow as suggested by (Pieh & Laufer 2006). Reproduction of released *T. scripta* has not been reported in southern Europe (Cadi *et al*. 2004; Perez□Santigosa, Díaz-Paniagua & Hidalgo□Vila 2008; Sperone *et al*. 2010; Sancho & Lacomba 2016; Foglini & Salvi 2017), but also in a part of Croatia with continental climate (Koren et al. 2018), as well as in central Europe in Slovenia (Standfuss et al. 2016). The regular reproduction of *T. scripta* reported here for Germany adds to previous records of successful reproduction in temperate climate.

Even though successful reproduction of *T. scripta* in Europe has been observed, it has been argued that this will not make the species invasive, as sex determination in this species is temperature dependent (Geiger & Waitzmann 1996). At low temperatures, only male hatchlings occur (Cadi et al. 2004). However, female hatchlings occurred both in Spain (Perez□Santigosa, Díaz-Paniagua & Hidalgo□Vila 2008) and in southern France (Cadi et al. 2004). The sex ratio of clutches is influence by the soil temperature, and both sexes can be produced from incubation temperatures between 28.3 and 30.6°C (Cadi et al. 2004). So far, no soil temperature data are available for Kehl, but air temperatures above 33°C during the day and above 20°C at night are common in summer and often occur for periods of 2 weeks and more. The hatchlings captured in the current study were too small to be sexed. Climate warming will make it more likely that females will hatch in the future.

To make management decisions for the long-living and slowly reproducing *T. scripta*, it is essential to estimate how these populations will develop for several decades. Most *T. scripta* that were sold in pet shops were females, as breeders in the USA used high temperatures for faster hatching, and higher temperatures induced a female biased sex-ratio (Prevot et al. 2007). The combination of a potential female biased sex ratio of released sliders (Prevot et al. 2007) and a male biased sex ratio of hatchlings could cause an increased population growth in the mid-term (5-20 years), as more females will lay fertilized eggs. An overall increase in the number of hatchlings together with rising temperatures due to global warming makes it more likely that female hatchlings will occur in the coming decades. This scenario makes it possible that *T. scripta* will establish viable populations in Kehl and other areas of the Upper Rhine Valley.

*Pseudemys nelsoni, P. concinna, Graptemys pseudogeographica* and *Mauremys sinensis* have a low risk of getting invasive in Europe (Kopecký, Kalous & Patoka 2013; Masin 2014). In contrast, *Trachemys scripta* is generally regarded as being invasive, as in many countries it can survive, breed, and colonise additional habitats with negative impact on the native biodiversity (Lowe et al. 2000). However, detailed studies on its effect on the native fauna and flora are rare. In France, adult *T. scripta* are omnivores, eating invertebrates, amphibians and fish, while older individuals have a more herbivore diet (Prevot et al. 2007). They have been observed to disturb the nests of water birds when searching for basking spots (Laufer 2007). They might have a negative impact on the endangered native European pond turtles (*Emys orbicularis*) by trying to mate with them (Jablonski et al. 2017) and by competing for the best basking spots with them (Cadi & Joly 2003). In Kehl, the Altrhein does not have an important native biodiversity, but there are ecologically important areas close by, especially the Rhien floodplains. Increasing population size at the Altrhein increases the risk that these habitats would be colonized. Further, *T. scripta* have also been observed next to the Kehler nature reserve Sundheimer Grund, an ecologically important and vulnerable habitat (Schradin, pers. observ.).

## Results with regards to German and European legislation

The regulation (EU) No 1143/2014 demands all member states to take actions against invasive species (European_Parliament & Council_of_the_European_Union 2014) including *T. scripta* (European_Commission 2016). After the trading ban in 2006 it was expected that the release of *T. scripta* into the wild will decline in Germany, and that the population in Kehl will decline due to mortality (Pieh & Laufer 2006). My current study 13 years later indicates that these assumptions were wrong. In Germany, it has been believed that, apart from informing the public not to release exotic pond turtles into the wild, no further actions are needed. It was argued that it is too cold for the released individuals to survive for long periods and to reproduce (Geiger & Waitzmann 1996; Pieh & Laufer 2006; Laufer 2007; Nehring 2016). Thus, populations in Germany are still officially believed to be unstable (Nehring 2016; StA_"Arten-_und_Biotopschutz” 2018b), which has influenced the national action plan (StA_"Arten-_und_Biotopschutz” 2018a). My study shows that these assumptions are wrong: Firstly, population size is not decreasing and there is no indication for high mortality. Secondly, successful reproduction occurs more regularly than previously believed. Thus, the management plan in Germany should be reconsidered.

The results presented here add to several other studies indicating that *T. scripta* has the potential to become invasive in western, middle and central Europe (Mačát & Jablonski 2016; Standfuss et al. 2016; Koren et al. 2018). Thus, action plans to avoid an invasion might not only have to be changed for Germany, but for many other European countries. For France, it has been suggested to remove sliders from all wetlands (Cadi et al. 2004), and this has been done in Spain (Sancho & Lacomba 2016). While the invasion is slower in colder regions, the fact that the exotic populations there don’t decline but reproduce and increase should be of concern.

## Conclusions

The population of exotic pond turtles at the Althrein of Kehl did not decreased since the ban of trade with *T. scripta* in the European Union. On the contrary, the diversity of species being released has increased. In addition, the population size of *T. scripta* is also not decreasing but seems to increase, and there is no indication of high mortality due to cold winters, but of regular successful reproduction in summer. This might lead to a more balanced sex ratio in the future and as such a further increase in reproduction and population size. It is now important to measure soil temperatures, to determine the sex ratio of hatchlings, and to combine this with climate simulations for the next 30 years. In sum, the current action plan in Germany for *T. scripta* will have to change if an invasion is to be avoided. Such an action plan could include (see also (Teillac-Deschamps et al. 2009) 1. Educating the public, 2. prosecute people releasing exotic pond turtles, for example by publishing photos of released pond turtles and searching for its owner, 3. destruction of clutches, 4. removal of large females from the population when they are been found on land before / after laying clutches, and 5. active removal of released exotic pond turtles, especially from ecologically valuable waters. Other west and central European countries might have to apply similar action plans.

## ACKNOWLEDGEMENTS

This project was funded by the CNRS. The project was done as part of a course I do at the Hector Kinderakademie in Kehl, teaching 8 to 9 years old children about animal behaviour, population biology, animal welfare and nature conservation. I am very thankful to the Hector Kinderakademie in Kehl for their support. I am thankful to members of the facebook group “Wasserschildkröten” which assisted me in identifying species from photographs, especially to N. Ziegenhagen, who runs an animal shelter for exotic pond turtles. I would also like to thank the city of Kehl (Dr. A.-M. Amui-Vedel, Bereich Stadtplanung/Umwelt) and the Regierungspräsidum Freiburg (S. Person) for their support for this study.

